# A model of transcriptional bursting dynamics based on coupling between chromatin and enhancer states

**DOI:** 10.1101/2025.11.29.691276

**Authors:** Ofri Axelrod, Udi Binshtok, Brian Gebelein, Hernan G. Garcia, David Sprinzak

## Abstract

In recent years, it has become evident that transcription is not always a continuous process. Rather, many genes exhibit bursting behavior characterized by discrete periods of transcriptional activity and inactivity. While transcriptional bursting has been broadly observed across different organisms, from bacteria to mammals, we lack a mechanistic understanding of the molecular events that regulate this widespread process. Specifically, how the expression of a ‘bursty’ gene is quantitatively determined by different molecular factors such as the concentration of transcription factors (TF) and architecture of the enhancers is not well understood. Here, we introduce a model based on the interplay between chromatin state and TF binding in order to describe bursting dynamics. We leverage the widespread Monod-Wyman-Changeux two-state model to predict the dependence of transcriptional bursting dynamics on TF concentration, binding affinity, and number of TF binding sites. We use the model to qualitatively reproduce the behavior of bursting dynamics observed in the *C. elegans* gonad. Overall, we provide a tractable model for transcriptional bursting that offers mechanistic insights into the factors regulating transcriptional bursting and generates experimentally testable predictions capable of uncovering the molecular basis of this widespread process.

## Introduction

As the mediator between genes and their protein products, transcription is a key process underlying the function of the cell. A few decades ago, McKnight and Miller showed hints of transcription being a discontinuous process. Here, steady transcription may be interrupted by periods of silenced expression, despite the constant molecular constituents within the cell^1^. In recent years, single molecule and live imaging methods, such as single molecule FISH^2^ and MS2/MCP-GFP tagging^3^, have been developed to track mRNA at transcription-initiation sites. These approaches allowed detailed scrutiny of transcription dynamics, confirming that, in many cases, transcription exhibits bursting behavior. This behavior is characterized by sharp switching between discrete periods of transcript production and periods without any transcription, termed ‘ON’ and ‘OFF’ transcriptional states (reviewed in Leyes et al^4^, Rodriguez & Larson^5^).

The simplest model that can be used to explain the bursting phenomenology is a two-state model. Here, a gene switches between active and inactive states, termed ‘ON’ and ‘OFF’ transcriptional states, whereby transcription is initiated only in the active state^6–9^. In this context, transcriptional bursts are characterized by three parameters: (i) the burst duration, which describes the mean period transcription is active (ON time), (ii) the burst separation, which is the mean time between bursts (OFF time), (iii) and the burst amplitude, indicating the rate of mRNA production during a burst. Many studies have assessed how different cellular and molecular factors affect bursting parameters (reviewed in Leyes et al^3^). For example, TF concentration and TF binding affinity regulate bursting dynamics in cell culture^8,10,11^ and in vivo^12–14^. Lee et. al. showed that decreased activation of Notch signaling (less activator) in the *C. elegans* gonad leads to shorter burst duration of a Notch target gene^12^, but the burst amplitude was largely unaffected by the magnitude of Notch activation. Similarly, a study in *Drosophila* showed that burst duration increased with Notch activity^13^. Other studies have linked the chromatin state and nucleosome binding to transcriptional bursting^15,16^. For example, in yeast, nucleosomes affect Gal4 dwell time on the promoter and consequently reduce bursting amplitude^17^. The association of chromatin state with transcriptional bursting also stems from the relatively long timescales observed for transcriptional bursting, which can range from minutes to hours, depending on the organism^12,18,19^. These timescales correlate with timescales observed for chromatin accessibility, nucleosome turnover, and histone modifications^20,21^. Further, recent single molecule imaging in yeast and mammals have associated the dynamics of bursting with cooperative binding to multiple TF binding sites^22^ and to the complex co-assembly of TF condensates, TF binding sites at the enhancer, and the promoter^23^. Hence, copious amounts of experimental evidence link the dynamics of transcriptional bursting to both TF binding dynamics and chromatin state.

Mathematical implementations of two-state models are generally described by transition rates from active to inactive states that control ON and OFF time dynamics^4,16^. These phenomenological models, while useful, typically lack a mechanistic connection between TF concentration, enhancer/promoter architecture (e.g. the number, position and affinity of TF binding sites), and bursting dynamics. Thus, current models cannot account for the dependence of bursting dynamics on TF concentration and enhancer properties.

To account for this gap, we consider a mechanistic two-state feedback model in which TF binding to multiple DNA sites is coupled to chromatin states (referred to as open and closed chromatin states). The model is mathematically equivalent to the widespread Monod-Wyman-Changeux^24^ (MWC) two-state model^25^ and has been previously used by Mirny^26^ to describe the average, steady-state response in a highly cooperative non-linear transcriptional system as a function of TF concentration. We expand the utility of the model suggested by Mirny to account for the unique temporal dynamics of transcriptional bursting. In this model, TF binding is coupled to closed and open chromatin states in two ways: (i) the affinity of TF binding to the enhancer in the closed chromatin state is lower compared to its affinity in the open chromatin state, and (ii) the transition rate from open to closed chromatin state decreases with increased TF occupancy^27^. Therefore, the chromatin state of the system affects TF occupancy on DNA, and, in turn, TF occupancy (which depends on TF concentration) feeds back into chromatin by affecting the transition times between the two chromatin states. The model is simple and allows systematic analysis of bursting dynamics with respect to relevant parameters and time scales of transcription. We apply this model to assess how bursting ON and OFF times change with TF concentration, the number and affinity of TF binding sites, and parameters associated with chromatin dynamics. Using existing data we show that the model can qualitatively account for the dependency of transcriptional bursting dynamics on TF concentration observed experimentally, as well as generate new testable predictions. Thus, our feedback model offers a concrete example of the type of dialogue between theory and experiment that can shed light on the molecular mechanics underlying transcriptional bursting.

## Results

### A two-state feedback model for transcriptional bursting

To theoretically describe transcriptional bursting dynamics, we developed a two-state feedback (TSF) model that couples TF binding and chromatin states. This coupling allows establishing direct relation between busting dynamics, controlled here by switching between chromatin states, and TF concentration. In our model, we consider an enhancer that can be found in two states termed closed and open chromatin states (Fig. 1A). The enhancer contains *N* binding sites for a TF controlling the transcriptional response of a gene. We note that the two states do not necessarily have to be associated with open and closed chromatin states but can describe any two transcriptional states that satisfy the assumptions of the model (e.g. condensed vs non-condensed DNA region, conformational shifts in transcription activation domains, etc.). We define two variables associated with the state of the system: (i) the occupancy, *n*, which describes the number of TFs bound to the enhancer out of its *N* total binding sites, and (ii) the chromatin state, *m*, defined by

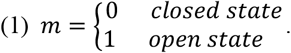

**Figure 1:**
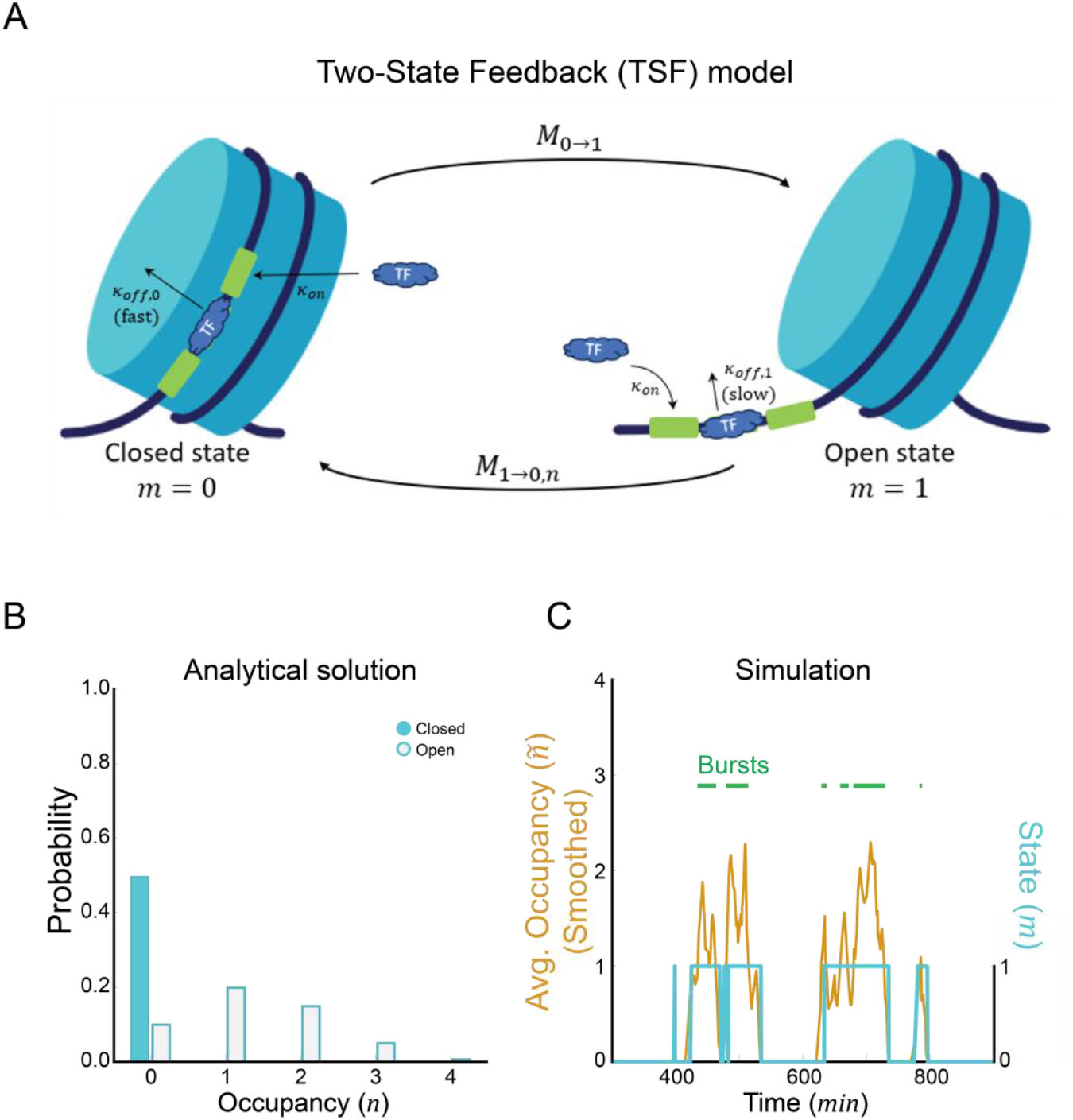
Analytical and numerical solutions of the TSF model show distinct occupancy distributions and transcriptional bursting dynamics. **(A)** Schematic of the TSF model. An enhancer with *N* binding sites (green boxes) can be in a closed (***m* = 0**) or open (***m* = 1**) chromatin state. Transcription factors (blue) can bind (with rate ***k***_***on***_) and unbind (with rate ***k***_***off***,***m***_) the binding sites. The unbinding rates, ***k***_***off***,***m***_, depend on the chromatin state (*m*). The transition between states is controlled by the rates ***M***_**0→1**_ and the ***M***_**1→0*n***_, with the latter depending on the number of occupied TF binding sites, *n*. Cartoon adapted from *BioRender*, prior to further editing. **(B)** Probability distribution of binding site occupancy at each of the chromatin states (Blue, ***p***_***n***,**0**_, ***Closed***. Orange, ***p***_***n***,**1**_, ***Open***), obtained from the analytical steady state solution. The overall probability of the system being in each of the chromatin states is shown in supplemental Fig. S1A. **(C)** Average occupancy (Blue; ***ñ*(*t*)**) and chromatin state (Orange; ***m*(*t*)**) dynamics from a single simulation (Example from time interval ***t*** ∈ [3**00**, 9**00**] ***min***). The average occupancy, ***ñ***, is obtained by time-averaging the occupancy using a moving window of 10 min (see methods and supplemental Fig. S1C-D). We define bursts as continuous time intervals where the occupancy is ***ñ* ≥ 1** (Examples of bursts are under the green lines). In panels **(B-C)** default parameters were used (Supplementary Table 1).

We assume simple binding kinetics of TFs to the enhancer. The binding rate of a single TF to a single DNA site is defined as *k*_*on*_. Here, *k*_*on*_ = *k*^+^[*TF*], where [*TF*] is the concentration of the TF in the nucleus and *k*^+^ is the first-order association rate of the TF to the binding site. The unbinding rate of the same TF from an occupied binding site is defined as *k*_*off,m*_, where the *m* subscripts indicate different unbinding rates for different chromatin states. Based on Donovan et al^16^ and Luo et al^28^, we assume that the chromatin state affects binding affinity such that *k*_*off*,0_ ≫ *k*_*off*,1_, where *k*_*off*,0_ is the unbinding rate in the closed state and *k*_*off*,1_ is the unbinding rate in the open state. This assumption introduces dependency of the binding/unbinding dynamics on the chromatin state of the enhancer. We do note that the binding rate, *k*_*on*_, may also depend on the chromatin state, but for simplicity, we consider here only the unbinding rate, *k*_*off*_, to be affected by the chromatin state (a model where *k*_*on*_ would depend on chromatin state would be mathematically equivalent).

When there are multiple binding sites in an enhancer, we assume that TF binding to each site is independent (i.e., there is no cooperativity). In this model, the binding rate of a single TF *K*_*on,n*_ Ito the enhancer is proportional to the number of free binding sites (*N* − *n*) such that

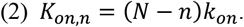

and the unbinding rate of any of the *n* bound transcription factors will be given by

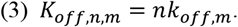

Finally, we assume that the chromatin can switch between open and closed states at some baseline rates. We define *M*_0→1_ as the transition rate from the closed to the open state and *M*_1→0,*n*_ as the transition rate from the open to the closed state, which depends on occupancy *n* (as described below (Eq. 9)). This assumption introduces dependency of the enhancer chromatin state on the TF occupancy of the enhancer. Specifically, this dependence enhances the probability of the chromatin to be in the open state when TFs are bound by decreasing the transition rate from the open to the closed chromatin state. Together, the dependency of TF dissociation rate on chromatin state and the dependency of chromatin state on enhance occupancy form a coupling between the chromatin state and the binding/unbinding dynamics of the TF. As we will see below, this coupling allows for the direct modulation of transcriptional bursting by changing TF concentration and the architecture of the enhancer.

The overall map of states, occupancy and transition rates is then defined by

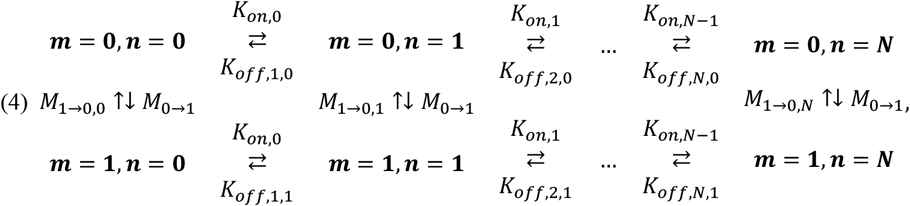

where *m* = 1 and *m* = 0 corresponds to the open and closed chromatin states, respectively, and *n* is the number of occupied TF binding sites. The dynamics of the system can be formalized as a set of master equations that describe the probabilities *p*_*n,m*_ of the system to be at a state *m* and occupancy *n*. As a result, the probabilities need to be normalized such that

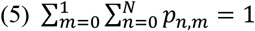

These probabilities can change in time through the unbinding or binding of TFs (moving between columns in eq. 4) or through a transition between chromatin states (moving between rows in eq. 4). Thus, the change in probabilities over time, *t*, is described by the set of 2*N* differential master equations given by^7^

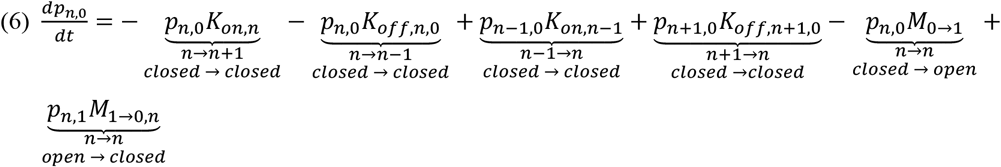

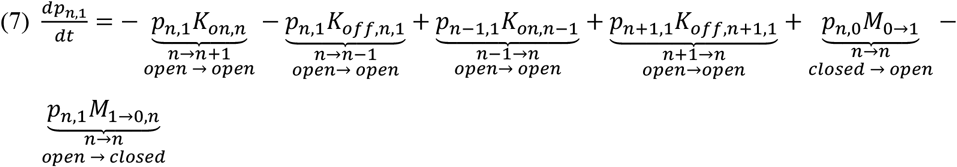

where we have indicated the state transition described by each term in the equations underneath the term. Assuming equilibrium conditions, the steady state solutions 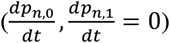 for these equations can be solved and the average occupancies can be calculated analytically^26^. A similar model was used by Mirny^26^ to show that the average occupancy and average chromatin state exhibit a non-linear cooperative response as a function of TF concentration. However, Mirny’s model was developed and solved solely in *steady-state*, and it did not consider the system’s dynamics. Here, we used our new formulation (eq. 6,7) to dissect the dynamics of the system as a function of the different parameters of the model.

In general, the system described here can operate either in equilibrium or outside equilibrium. The requirement for equilibrium is not necessarily valid for many biological systems, as energy is often invested in processes such as histone modification (e.g. methylation and acetylation) associated with chromatin states^29^. Nevertheless, we choose to consider an equilibrium condition, as this is the simplest assumption that can be made for the system, and thereby allows for a focused exploration of the predictions of transcriptional bursting dynamics stemming from this two-state model.

To enforce equilibrium, we invoke the requirement for detailed balance^30^. Namely, that the probability flux around any closed loop along the state map in one direction is equal to the probability flux corresponding to moving in the opposite direction. Mathematically, this can be shown to be the same as demanding that the product of the rates in one direction be equal to the product of the rates in the opposite direction^30^. Starting in the top left corner of eq. 4 (*m* = 0, *n* = 0), the rates along any closed loop that goes clockwise to some occupancy *n* along the top row in eq. 4 and then goes back through the bottom row of eq. 4 should be equal to the corresponding rates along the same loop going counterclockwise. This requirement results in the following relations

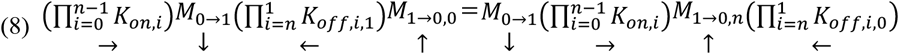

Here, the arrow below each term indicates the direction of rates along the clockwise and counterclockwise loops in Eq. 4, going from zero occupancy to some *n* occupancy and back. Simplifying and plugging in equation (3) into equation (8) (See Appendix) leads to the relation between closing rates at general occupancy *n* and at *n* = 0 given by

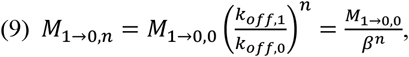

where 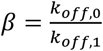 is the fold change in the TF unbinding rate between the closed and open chromatin states. Thus, since we assumed that *β* ≫ 1 (*k*_*off*,0_ ≫ *k*_*off*,1_), the requirement of equilibrium leads to an exponential decrease in the chromatin closing rate with increased occupancy, namely, the presence of bound TFs promotes the open state.

An earlier two-state model by Lammers et al. assumed no detailed balance, and therefore no dependence of closing rate on occupancy. To generalize the TSF model and allow comparisons with the previous model, we define the dependence on occupancy as

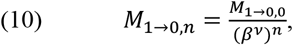

where *v* = 1 corresponds to the equilibrium case, while *v* = 0 represents the non-equilibrium situation where opening and closing rates do not depend on occupancy (as in Lammers et al. 2020).

To identify the key parameters in the TSF model, we defined a set of dimensionless variables that capture the relations between the different time scales of the model given by

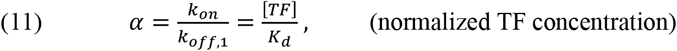

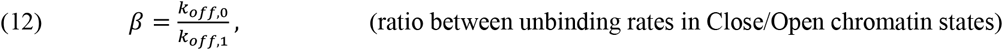

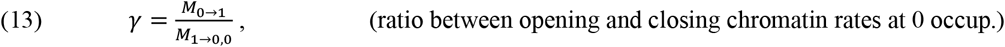

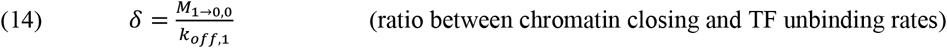

where 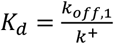 is the equilibrium dissociation constant of a TF to a single DNA binding site when the chromatin is open. We also scale the time by the unbinding rate of TFs from a single binding site in the open state, namely

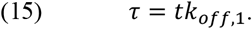

Using these dimensionless parameters in (11) to (15), equations (6) and (7) become:

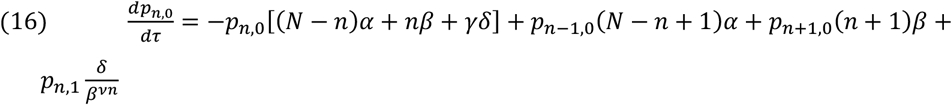

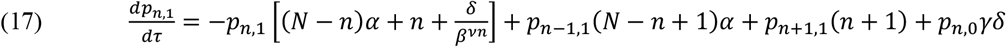

In the coming sections, we analyze equations (16) and (17) and obtain the probability distributions describing the chromatin state *m* and occupancy *n*, as well as the dynamics of transitions between states. We test the effect of changing the TF concentration (represented by *α*), the number of binding sites, *N*, the dimensionless parameters *β, γ, δ*, and the factor *v* on our theoretical predictions to generate testable experimental predictions. In doing so, we provide a framework to explore and test molecular factors of transcriptional bursting.

### The TSF model exhibits state-dependent occupancy distributions leading to transcriptional bursting

We first sought to determine whether the model can recapitulate transcriptional bursting, switching between active and inactive periods of transcription in a given parameter regime. We adopted the “occupancy hypothesis” which assumes that transcription rate is proportional to the average occupancy of TFs on the enhancer, an assumption that has been recently confirmed experimentally in mammalian cells^16,21,27^. In this context, it is expected that bursting behavior will be characterized by distinct occupancy distributions for each chromatin state, corresponding to ON and OFF transcriptional states. We first considered a set of default parameters determined based on previous work^16^ and reasonable biological constraints (Supplementary Table 1). We set the number of binding sites, *N*, to 4. Further the *α* parameter was set to 0.5, which corresponds to a situation where the concentration [*TF*] is equal to half the dissociation constant *K*_*d*_. As a reminder, we assume that the unbinding rate of TFs from a single binding site is much faster in the closed chromatin state than in the open chromatin state (i.e., *β* ≫ 1). We also assume that, when none of the sites are occupied (*n* = 0), the transition to the closed state is faster than the transition to the open state (that is *γ* < 1). Finally, since chromatin dynamics are slower than TF dynamics (molecular timescales are summarized comprehensively in Lammers et al^16^), we assume that the transition between chromatin states is slower than TF unbinding (that is *δ* < 1).

Following these assumptions, equations (16) and (17) were solved analytically in *steady-state* and the solutions of the probabilities *p*_*n*,0_ and *p*_*n*,1_ were visualized as a function of *n* (Fig. 1B). We use these expressions to define the probability of the system being in the closed state, *S*_0_, or in the open state, *S*_1_ (Supplemental Fig. S1A), which are given by

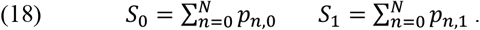

Under these assumptions, the *steady-state* solution shows two distinct occupancy distributions with two distinct peaks. One peak is centered at higher occupancy levels and is associated with the open state (orange bars in Fig. 1B). The second peak is centered at zero occupancy and is associated with the closed state (blue bars in Fig. 1B). This result is consistent with transcriptional bursting, where the system dynamically alternates between transcriptional regimes of transcription ON and transcription OFF at a constant TF concentration. We note that the occupancy distribution obtained from long term simulation recapitulates the analytical solution (Supplemental Fig. S1A,B).

To test if the TSF model displays dynamics of transcriptional bursting, we performed numerical simulations that calculate the chromatin state and occupancy of the system at every time point, over extended periods of time, using the Gillespie algorithm^31^ (see Methods). As described above, we assumed that the transcriptional activity from a promoter controlled by the enhancer is proportional to the average TF occupancy on the enhancer over a fixed time scale^16,21,27^. Since transcriptional events typically involve multiple biochemical processes (polymerase recruitment, initiation, elongation) that occur on a longer time scale compared with TF binding and unbinding dynamics (minutes vs seconds)^32^, we smoothed the occupancy dynamics with a moving time window of 10 minutes (see Supplemental Fig. S1C-D and Methods). Note that this smoothed occupancy, defined as *ñ*, results in non-integer values for the average occupancy (Supplemental Fig. S1D). The simulations of the average occupancy with the default parameters indeed exhibited bursting behavior, switching stochastically between high and low occupancy states at long molecular timescales (tens of minutes, Fig. 1C). To obtain the distributions of the mean durations of active and inactive transcription (ON and OFF times), we defined a transcription ‘ON’ state when *ñ* ≥ 1 (marked by green dashes in Fig 1C), and a transcription ‘OFF’ state when *ñ* < 1. Overall, the TSF model exhibits bursting dynamics, reflected both in the occupancy distribution and the temporal analysis.

### Dynamics of transcriptional bursting depend on transcription factor concentration

Next, we tested how the occupancy probability distributions, and corresponding bursting dynamics, are affected by changes in TF concentration. Therefore, we varied the parameter *α*, accounting for the normalized TF concentration, and tracked both the occupancy probability distributions and smoothened occupancy of TF binding to the enhancer (Fig. 2). All other parameters were held at fixed default values (Supplementary Table 1). We found that at low *α* values the system is mostly in the closed state, with very short bursts of transcription (Fig. 2A, B). As the value of *α* increases, there is a higher probability for the system to be in the open state and bursts become prolonged (Fig. 2A’, B’). When values of *α* are of the order of 1 and higher, the system is primarily in the open state, yet there are relapses to the closed state leading to a bimodal occupancy distribution (Fig. 2A’’, B’’).

**Figure 2:**
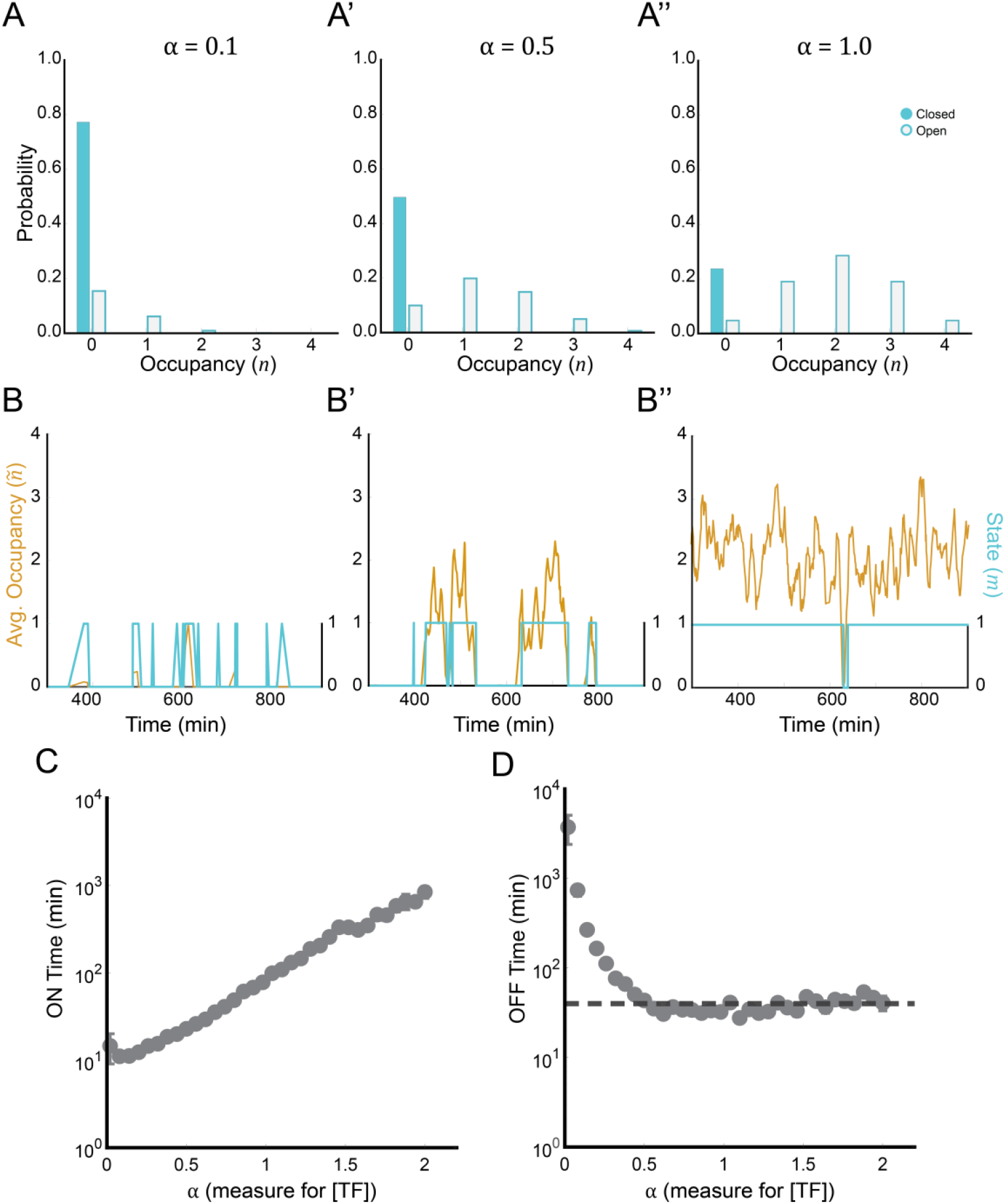
Dependence of bursting dynamics on TF concentration in the TSF model. **(A-B’’)** Probability distributions (A-A’’) and corresponding average occupancy dynamics (B-B’’) for different values of 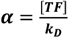 (as indicated). **(C-D)** Dependence of ON time (C) and OFF times (D) on the value of ***α***. Each data point is an average ON time and average OFF time, calculated by analysis of a simulation ran over 30,000 minutes and taking the average of continuous time periods when ***ñ* ≥ 1** (ON) and average of continuous time periods when ***ñ*** < **1** (OFF). Error bars are the standard error of the mean (SEM). In all panels, default parameters were used and only ***α*** was changed as indicated (Supplementary Table 1).

We next use this analysis to track how the average ON and OFF times change with TF concentration, or *α* in our model. This analysis holds biological importance, as the ON and OFF times can be measured experimentally. We find that the average ON time increases exponentially with *α* while the average OFF time initially decreases with *α*, and then reaches a baseline value (Fig. 2C, D). The baseline value of the OFF time is approximately equal to the inverse of the opening rate, *M*_0→1_. We note that since *α* is the ratio between TF concentration and binding affinity, this analysis is relevant for changing either of these parameters. Thus, the TSF model provides a quantitative description for how bursting dynamics change with TF concentration and binding affinity, and shows how bursting behavior persists even at high TF concentration. Moreover, the model suggests regimes in which increasing TF concentration will not affect the OFF time but will increase the ON time (*α* > 0.5 in Fig. 2C,D), a result consistent with previously shown experimental results, as discussed in discussed in the sections below. We note that effects on the amplitude of bursts were not analyzed since the bursting amplitude depends on the conversion from average occupancy to transcription initiation. This conversion may depend on multiple biological factors which are not directly represented in the model.

### Models without feedback do not exhibit essential features of transcriptional bursting

To understand the importance of feedback between occupancy and chromatin state, we considered parameter limits without any feedback mechanism. In the TSF model, we can decouple chromatin state and occupancy in two ways. One way is to set the parameter *v* to 0 (resulting in the model considered in Lammers et al., 2020). In this case, the transition rate from open to closed states is fixed at its maximum (*M*_1→0,0_) and does not depend on the occupancy state, *n*. In this model, the occupancy probability distribution still depends on chromatin state. However, the probability of the system to be in the closed or open states is independent of TF concentration (Supplemental Fig. 2A). Hence, the probability distribution in the closed state remains effectively unchanged over an order of magnitude change in *α*. On the other hand, the probability distribution in the open state shifts to higher values of *n* as *α* was increased. Furthermore, the burst dynamics did not exhibit strong dependence on *α*, with ON times remaining at about the same level across a wide range of *α* values (Supplemental Fig. 2B-C). The OFF times do initially decrease with *α* (since more bursts emerge), but they only change moderately at high *α* values as the burst frequency remains the same (Supplemental Fig. 2D). Thus, increasing values of *α* in this model (*v* = 0) affects the occupancy level, but burst durations remain similar, as chromatin transition rates are independent of TF occupancy.

A second way to decouple the chromatin state and occupancy in the TSF model is to set the parameter *β* to 1 (i.e., *k*_*off*,0_ = *k*_*off*,1_). In this case, the dissociation rate of TFs from a single binding site is not modulated by the chromatin state *m*. Occupancy distributions of this model are state-independent, meaning that both the closed and open state distributions are similar. Further, these distributions shifted together to higher values of *n* as *α* increased (Supplemental Fig. 2E). The average occupancy level, *ñ*, still increases with *α*, but as *α* increases the system exhibits noisy continuous levels of transcription rather than bursting (Supplemental Fig. 2F). Since the average occupancy level in this model exhibits stochastic fluctuations with a similar distribution between chromatin states, the average ON time increased faster than in the TSF model (Supplemental Fig. 2G) and the OFF time decreased more quickly with *α* values compared to the TSF model (Supplemental Fig. 2H). Importantly, the OFF time decreases to zero when increasing *α*, which is uncharacteristic of transcriptional bursting.

Overall, these models show that MWC-like models of transcriptional bursting require feedback between the chromatin state and TF occupancy to exhibit bursting dynamics that depend on TF concentration.

### Transcriptional bursting dynamics depend on the number of TF binding sites

Since regulatory regions of genes vary in their number of TF binding sites^33,34^, we sought to characterize the effects of the total number of binding sites, *N*, on the transcriptional response generated by the TSF model. Moreover, TF binding site number is a parameter that can be easily manipulated experimentally, making it a prime target for generating actionable predictions using our model. We found that increasing values of *N*, without changing any other parameter, resulted in a higher probability for the system to be in the open state (higher *S*_1_). Furthermore, we found that the dependency of the occupancy probability distribution on the state becomes more pronounced, even reaching bimodality, as the distribution of open states moves to higher *n* values with increasing numbers of TF binding sites (Fig. 3A). Our simulations further show that there is an increase in ON times with increasing numbers of TF binding sites. However, the dependence of OFF times on *N* is weak (Fig. 3B,D). Overall, adding more sites increases the bimodal nature of the system and promotes an increase in occupancy levels and ON times.

**Figure 3:**
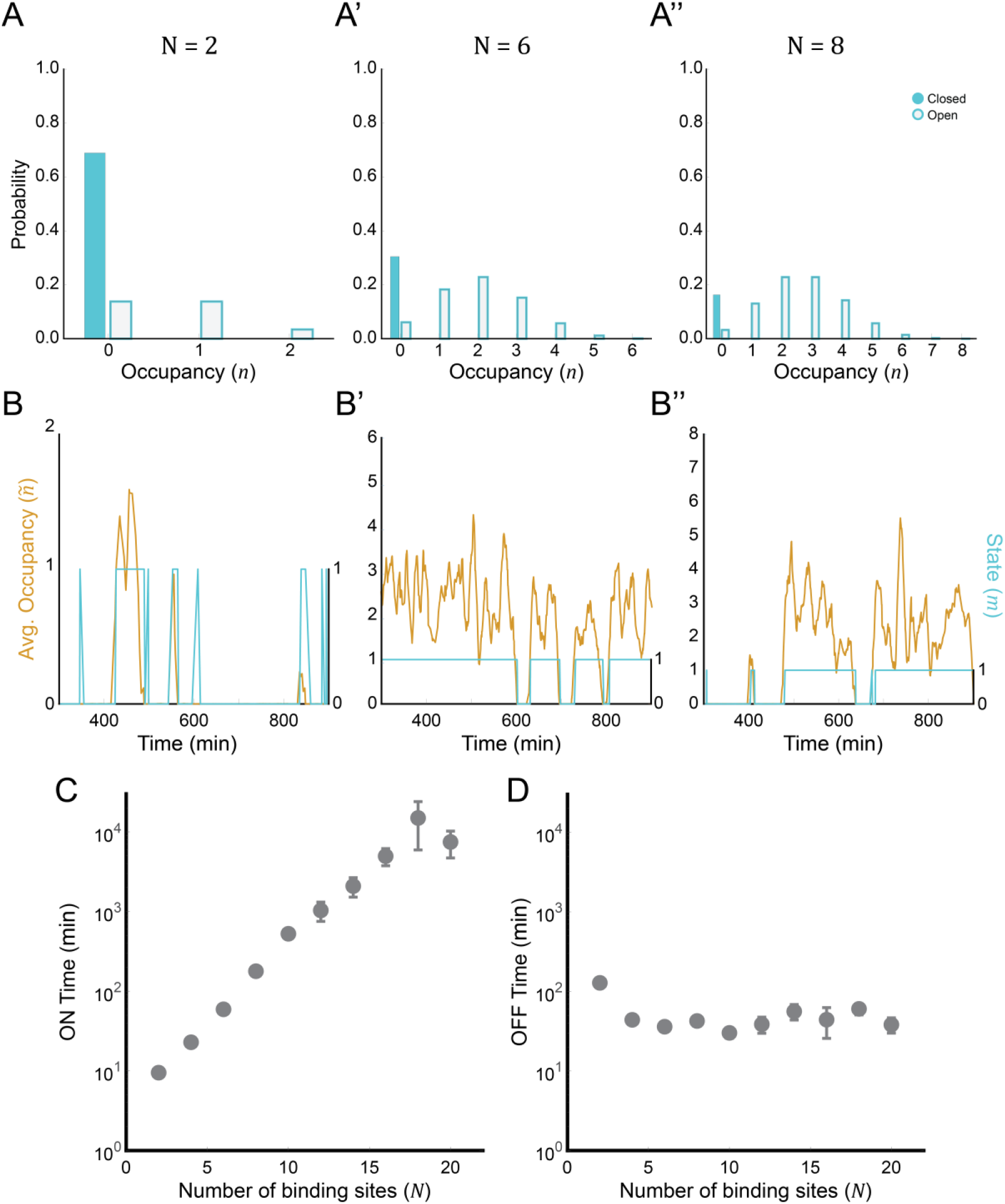
Increase in number of TF binding sites results in longer transcriptional bursts. **(A-B’’)** Probability distributions (A-A’’) and corresponding average occupancy dynamics (B-B’’) for different number of TF binding sites, N (as indicated). **(C-D)** Dependence of ON times (C) and OFF times (D) on N. Each data point is an average ON time and average OFF time, calculated by analysis of the entire 30,000 minutes simulation and taking the average of continuous time periods when ***ñ* ≥ 1** (ON) and average of continuous time periods when ***ñ*** < **1** (OFF). Error bars are the standard error of the mean (SEM). In all panels, default parameters were used and only ***N*** was changed as indicated (See also Supplementary Table 1).

### The TSF model is robust to changes in parameters associated with chromatin dynamics

So far, we have analyzed the effect of TF concentration/binding affinity (*α*), TF binding site number (*N*), and relative unbinding rate (*β*) on bursting dynamics (Figs. 2, 3, and Supplemental Fig. 2, respectively). We next sought to determine the magnitude by which chromatin dynamics affect bursting dynamics in our model. To do so, we analyzed how a change in the parameters of the system relating to chromatin states, *γ* and *δ*, affect the outcome of the TSF model.

The parameter 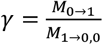 describes the ratio between opening and closing rates, which can shift the balance between the probability of the system to spend time in a closed versus open state (higher *γ* means a higher probability to be in the open state). Simulating the model at two values, *γ* = 0.1 and *γ* = 1, shows that at the higher *γ* the probability distribution is indeed shifted to a more open state, and that transcriptional bursts are more frequent (keeping all other parameters the same, Supplemental Fig. 3A, B). Simulating over a range of *α* values shows that the ON times are only weakly affected when *γ* is modulated, whereas OFF times decrease significantly. These results reflect the increased probability to be in the open state as *γ* increases (Supplemental Fig. 3C, D). Hence, the effect of increasing *γ* is to shift the baseline of the OFF times to lower values.

The parameter 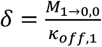 describes the relative time scale associated with chromatin closing rates and the TF unbinding rate (when the system is in the open state). Simulating the model at two values, *δ* = 0.1 and *δ* = 1, shows that in both cases, bursting occurs with similar dynamics (Supplemental Fig. 3E,F). Moreover, simulating over a range of *α* values shows that the ON and OFF times seem to be only mildly sensitive to this parameter, and only at higher *α* values (Supplemental Fig. 3G,H).

Overall, changes in both parameters do not alter the essential bursty behavior of the system, but rather change the quantitative response (ON and OFF times) to different TF concentrations.

### The TSF model recapitulates experimental observations

To validate the TSF model, we aimed to recapitulate observations from experiments directly measuring transcriptional bursting dynamics. We focused on a study by Lee et. al.^12^ that measured transcriptional bursting in the Notch target gene *sygl-1*, using the MS2/MCP-GFP tagging system in germline stem cells (GSCs) in the gonad of the nematode *C. elegans*. The maturation of the *C. elegans* gametes is coupled to a decreasing gradient of Notch activation, which functions as an activator (*α*) of the *sygl-1* gene. It has been shown that transcription of *sygl-1* in the GSC pool exhibits bursting dynamics, and that burst durations (ON times) is graded along the distal-proximal axis (Fig. 4A-B). At the distal end of the gonad, Notch signaling is highly active and *sygl-1* transcriptional bursting durations are the longest. As the distance of cells from the distal end increases, Notch signaling decreases. This causes the ON times to decrease monotonically in the GSCs as a function of their distance from the niche (along the distal-proximal axis; Fig. 4B). The OFF times were also analyzed for the same GSCs (Figure 4Fig. 4C). The experimental data was measured in two genetic backgrounds: in a wildtype (WT) condition and in a Notch receptor hypomorph mutant (*glp-1*(q224)) condition, which reduces the levels of Notch-dependent *sygl-1* expression. In GSCs at the same positions relative to the niche, ON times are decreased in the mutant compared with WT (Fig. 4B), while OFF times are increased (Fig. 4C). The nature of this experimental measurement, which is a recording of transcriptional bursting with an activation gradient, allowed us to test if our model can qualitatively capture the experimental results. Thus, we designed simulations of the TSF model assuming an enhancer with *N* = 4 binding sites (Supplementary Table 1), matching the 4 binding sites of the Notch activation complex in the *sygl-1* enhancer^35^. Since a quantitative concentration profile of the TF controlling *sygl-1* is unavailable, we assumed a decreasing sigmoidal function of *α* as a function of distance from the distal end, *x*, defined by 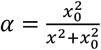 (Fig. 4D, left). With our default parameters, the simulations qualitatively recapitulated the decay profile in ON times (Fig. 4E) and the mild increase in OFF times (Fig. 4F) along the distal-proximal axis. To test if the TSF model can also capture the mutant condition, we performed the same simulations but reduced the level of *α* by a factor 2, namely, 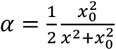 (Fig. 4D, right) as suggested by measurements reporting a corresponding decrease of Notch levels in the hypomorph animal (based on Lee et. al.^36^). Lowering *α* resulted in shorter ON times and weaker dependence of OFF times on the distance from the distal end (Fig. 4E,F), qualitatively matching the experimentally observed results. Overall, this analysis shows that the TSF model can capture qualitative features of experimental observation that did not have theoretical underpinning beforehand.

**Figure 4:**
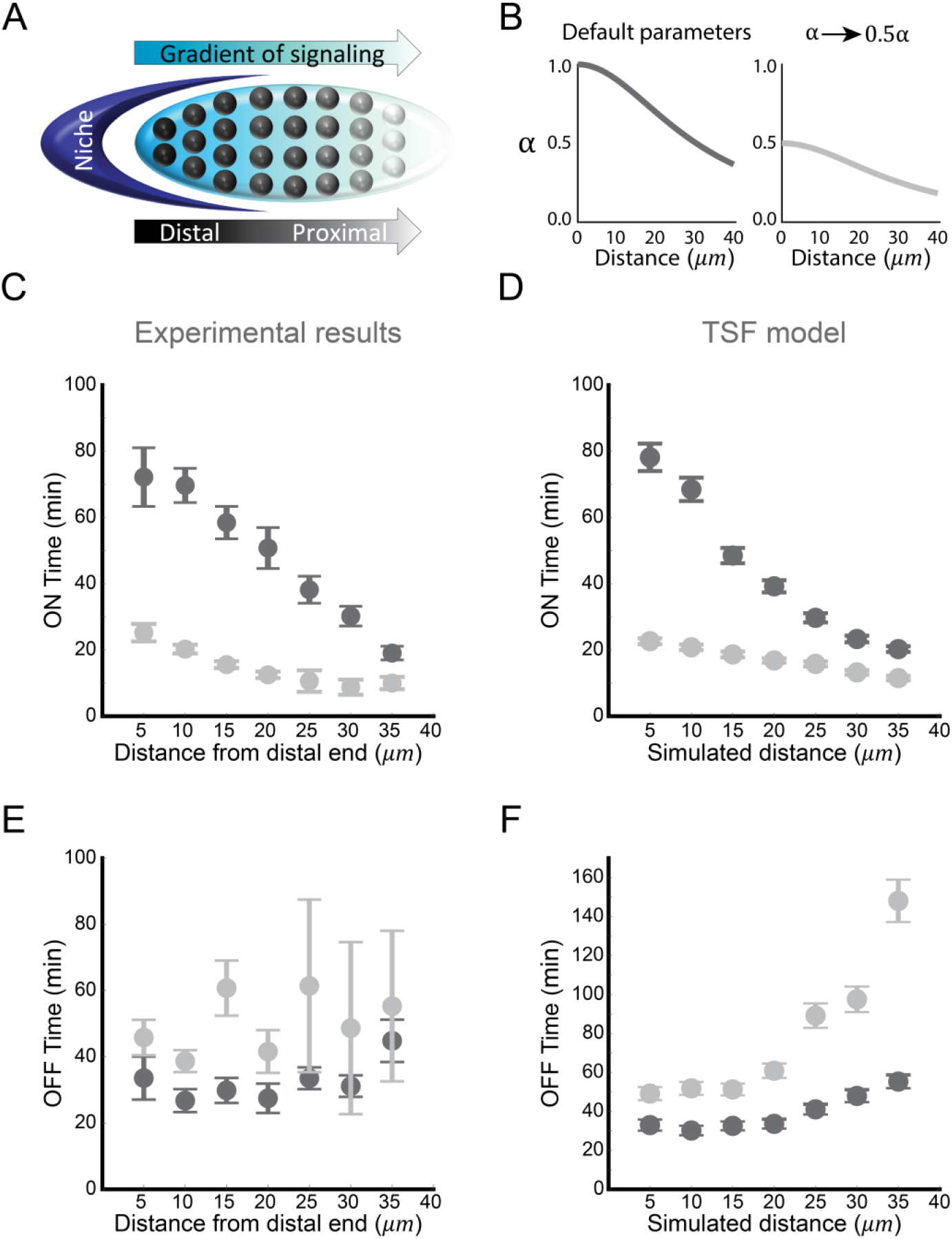
The TSF model recapitulates experimental results from a Notch target gene in the *C. elegan*s gonad. **(A-C)** Experimental results of Notch-dependent *sygl-*1 transcriptional bursting dynamics in germline stem cells (GSCs) of developing *C. elegance* gonad, taken from Lee et al.^**12**^. Schematic (A) shows the Position of GSCs, marked by circles, and Notch signaling gradient along the distal-proximal axis, relative to the niche. Experimentally determined ON times (B) decrease as a function of distance from the niche and with a hypomorph Notch mutant (*glp-1*(q224)). Corresponding OFF times (C) measured in same cells as in (B). **(D)** Spatial profiles of ***α*** assumed in the simulations. Sigmoidal decreasing profiles of ***α*** utilized were 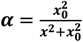 for WT Notch allele (left), with ***x*** being the position along the proximal-distal axis. 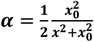 was used for the mutant Notch allele (right) corresponding to a reduction of Notch concentration by a factor of 2 in the mutant. ***x***_**0**_ was set to **30 µm. (E-F)** Simulations of ON times (E) and OFF times (F) for the WT and mutant *sygl-1* enhancer. We simulated the *sygl-1* enhancer by assuming ***N* = 4** binding sites. Each data point is the average ON time (E) and average OFF time (F), from a simulation over a time range of **[0, 30000**] minutes. Error bars are standard error of the mean (SEM). All parameters except ***α*** were set to the default values (Supplementary Table 1)

The TSF model is also consistent with the results of Falo-Sanjuan et al.^13^, who studied Notch-dependent transcriptional bursting during early embryonic development in *Drosophila*. In this work, the authors looked at the transcriptional bursting of the Notch target gene *single-minded* (*sim*) in the mesoderm and found that increasing Notch activity level led to a significant increase in ON times but almost no effect on OFF times. A similar trend, where the ON times increase and OFF times do not change with signaling strength, is observed in the analysis of target genes downstream of BMP signaling gradient in *Drosophila* embryos^37^. Our simulations show that there are broad parameter regimes where such qualitative responses are indeed expected (Fig. 2C,D, and Fig. 4E,F).

## Discussion

### Feedback mechanism supports transcriptional bursting

Transcriptional bursting has emerged as a ubiquitous feature of transcription observed across a plethora of organisms^4^. However, despite this ubiquity, we lack a mechanistic understanding of transcriptional bursting that functionally links bursting dynamics (ON and OFF times) with molecular details such as TF concentration and enhancer architecture. In this study, we aimed to provide a mechanistic model, based on the classical MWC^24^ and Mirny^26^ models that can account for the main features of transcriptional bursting, and generate predictions that can be tested experimentally. The model is based on coupling two activity-states: namely, chromatin opening/closing and TF binding dynamics to enhancers. The proposed TSF model incorporates parameter flexibility based on a two-way feedback. On the one hand, the chromatin state affects the binding affinity of TFs. On the other hand, the occupancy of enhancer binding sites by TF affects the transition from an open to closed chromatin state. The TSF model generates experimentally testable insights into how bursting dynamics depends on TF concentration and enhancer architecture (e.g. number and affinity of binding sites). In its basic form, the model relies on only four dimensionless parameters, thus allowing a comprehensive predictive analysis with relative ease.

### Dependence of transcription OFF and ON times on TF concentration and architecture

While TF concentration affects transcriptional bursting, it is unclear how bursting parameters are mechanistically associated with changes in TF concentration. The TSF model provides such a mechanistic relationship and generates testable predictions. The model suggests that increasing TF concentration leads to an exponential increase in the ON times. In contrast, the OFF times initially decrease with TF concentration but then plateau at a value inversely proportional to the baseline closing rate (equal to the minimal closing rate in the model). In other words, the TSF model predicts an increase in the duration of bursts, but not necessarily their frequency, as TF concentration level increases. We note that we did not consider in our model how bursting amplitude is affected by TF concentration and enhancer architecture since the bursting amplitude depends on the conversion from average occupancy to transcription initiation, which may depend on multiple factors not currently represented in the model.

Importantly, we showed that the TSF model can qualitatively match experimental results obtained in *C. elegans* (Fig. 4) and *D. melanogaster*^37^, where TF concentrations were varied. This finding is an indication of the strength of this model as a tool for generating testable predictions and accounting for observed experimental phenomena. A more quantitative fit to the model results will require specifically designed experiments where additional experimental details such as the TF concentration profile and binding affinity to the binding sites are controlled.

Our model also provides insights into the dependence of bursting dynamics on the number of binding sites in the enhancer region. Varying this parameter is often experimentally accessible, as binding sites can be mutated or introduced into reporter constructs with relative ease. Our model suggests that increasing the number of binding sites will lead to an increase in ON times while only mildly affect OFF times. Thus, we expect that experimentally increasing the number of binding sites in an enhancer will promote effects akin to increasing TF concentration. We note that the model can be used to assess changing the affinity of binding sites as well since all the model outputs depend on the ratio between the TF concentration and the affinity through the parameter *α*. Moreover, the model can be extended to consider enhancers with multiple sites having different affinities. The binding and unbinding rates in this case need to be accounted separately for each site.

### Feedback is required for bursting behavior

Our investigation of the model and its parameters showed that feedback is necessary for the manifestation of bursts and their dependency on TF levels. We show that a model where the transition between chromatin states does not depend on the occupancy states (*v* = 0, Lammers’ model) does exhibit bursting, albeit with fixed ON times that are independent of TF levels. Moreover, we show that a model where TF unbinding does not depend on chromatin states (*κ* = 1) fails to exhibit bursting dynamics. Thus, both ‘sides’ of the feedback are necessary for generating experimentally observed transcriptional bursting.

We note that the assumption that binding affinity depends on chromatin state is consistent with experimental observations showing that the dwell time of transcription factors on DNA is significantly shorter when a nucleosome occupies the binding sites^28,17^. Since *k*_*off*_ is defined as the rate of dissociation from the DNA (the inverse of dwell time), we assumed in our model that this unbinding rate, but not the binding rate (*k*_*on*_), is affected by chromatin state. Importantly, the model can be easily extended to the case where both binding and unbinding rates are affected by the chromatin state. This extension is expected to lead to a mathematically equivalent model and similar behavior of bursting dynamics.

### Expanding the model – cooperativity, repressors, and out-of-equilibrium behavior

The proposed TSF model can be expanded in a straightforward manner to include additional assumptions, depending on the system studied. For example, extended models may consider cooperative binding of TFs, by incorporating cooperativity (or anti-cooperativity) into the binding and unbinding rates^16,38^. An effect of cooperative binding sites on transcriptional bursting was observed for Notch target genes that contain paired sites promoting cooperative binding of Notch activators^13^. Another possible extension to the model is the incorporation of repressors, which can either bind to the same sites as the activators or different ones. Combinatorial transcriptional models have been used to generate different functional outputs^39^. While studies about the effect of repressors on transcriptional bursting dynamics are scarce, recent work suggest that repressors act by decreasing the frequency of transcriptional bursting by seemingly increasing the OFF time^40^.

A central feature of the current TSF model is that it is an equilibrium model. The requirement of equilibrium allows us to use the principle of detailed balance to define how the chromatin closing rate depends on TF occupancy and hence reduces the number of assumptions and parameters. Several recent works have suggested that equilibrium models may not be sufficient to explain transcriptional responses^29,41–43^. While non-equilibrium processes may well be important for transcriptional bursting, our model suggests that many of the experimentally observed features of transcriptional bursting can be captured by an equilibrium model. In the future, the TSF model can be easily extended to study the effect of relaxing the equilibrium assumption by exploring models that do not satisfy the detailed balance (as we have done for the case of *v* = 0). In that sense, our model can be a starting point for identifying features of transcriptional responses that require non-equilibrium processes.

## Methods

### Gillespie simulation

We used the Gillespie algorithm to simulate the dynamics of the system^31^. These simulations solve the dynamics of the system numerically by iteratively calculating the occupancy, *n*, and state of the system, *m*, over time. The parameters used are listed in Supplementary Table 1.

The iterations of the simulations operate as follow:

1. Considering a given state to the system (*n, m*), the rates for all possible transitions are calculated. The possible transitions are ‘bind’ (*n* → *n* + 1), ‘unbind’ (*n* → *n* − 1), and open/close (*m* = 0 → *m* = 1 *or m* = 1 → *m* = 0). These rates are determined according to:
  a. *rate*_*bind*_(*t*) = (*N* − *n*(*t*))*k*_*on*_
  b. *rate*_*unbind*_(*t*) = *n*(*t*)*k*_*off,m*_
  c. 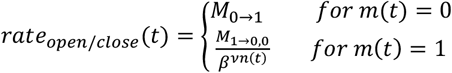
  d. *rate*_*total*_ (*t*) = *rate*_*bind*_(*t*) + *rate*_*unbind*_(*t*) + *rate*_*transition*_(*t*)
2. Calculating probabilities for events. The probability for each event to occur is calculated based on its rate and the sum of the rates of all possible events.
  a. 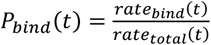
  b. 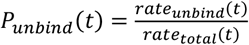
  c. 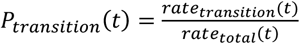
3. Determine and execute the next event. A number between 0-1 is chosen pseudo-randomly *P*_*uniform*_(*t*) ∈ *U*(0,1). An event, marking the next state of the system, is decided based on the outcome of the number and the probability for each event.
  a. If *P*_*uniform*_ ∈ (0, *P*_*bind*_] execute event “bind”
  b. If *P*_*uniform*_ ∈ (*P*_*bind*_, *P*_*bind*_ + *P*_*unbind*_] execute event “unbind”
  c. If *P*_*uniform*_ ∈ (*P*_*bind*_ + *P*_*unbind*_,1) execute event “bind”
4. Calculate the time step of the event and advance the simulation. The time for the event, *τ*(*t*), is chosen from an exponential distribution with mean 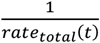. The new simulation time is updated to *t*_*new*_ = *t* + *τ*(*t*).
5. The new state of the system is updated based on the transition that has been chose.
6. Repeat steps 1-5.

### Parameters used to generate plots

Supplementary Table 1 below summarizes the parameters used in the analytical and numerical solutions of the TSF model, to generate the results shown in the main and supplementary figures. Most parameters were based on experimentally relevant ranges taken from Lammers et al^16^. The dissociation rate of the activator, *k*_*off*,1_, was based on a recent estimation of RBPJ/Notch TF residence time on its DNA sites^44^.

**Supplementary Table 1:**
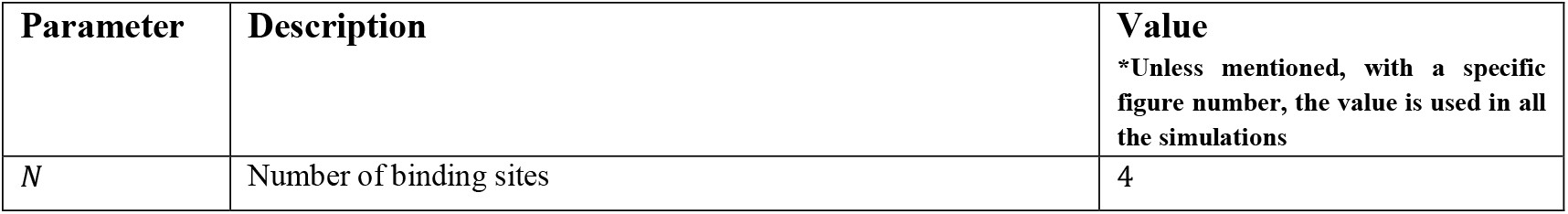
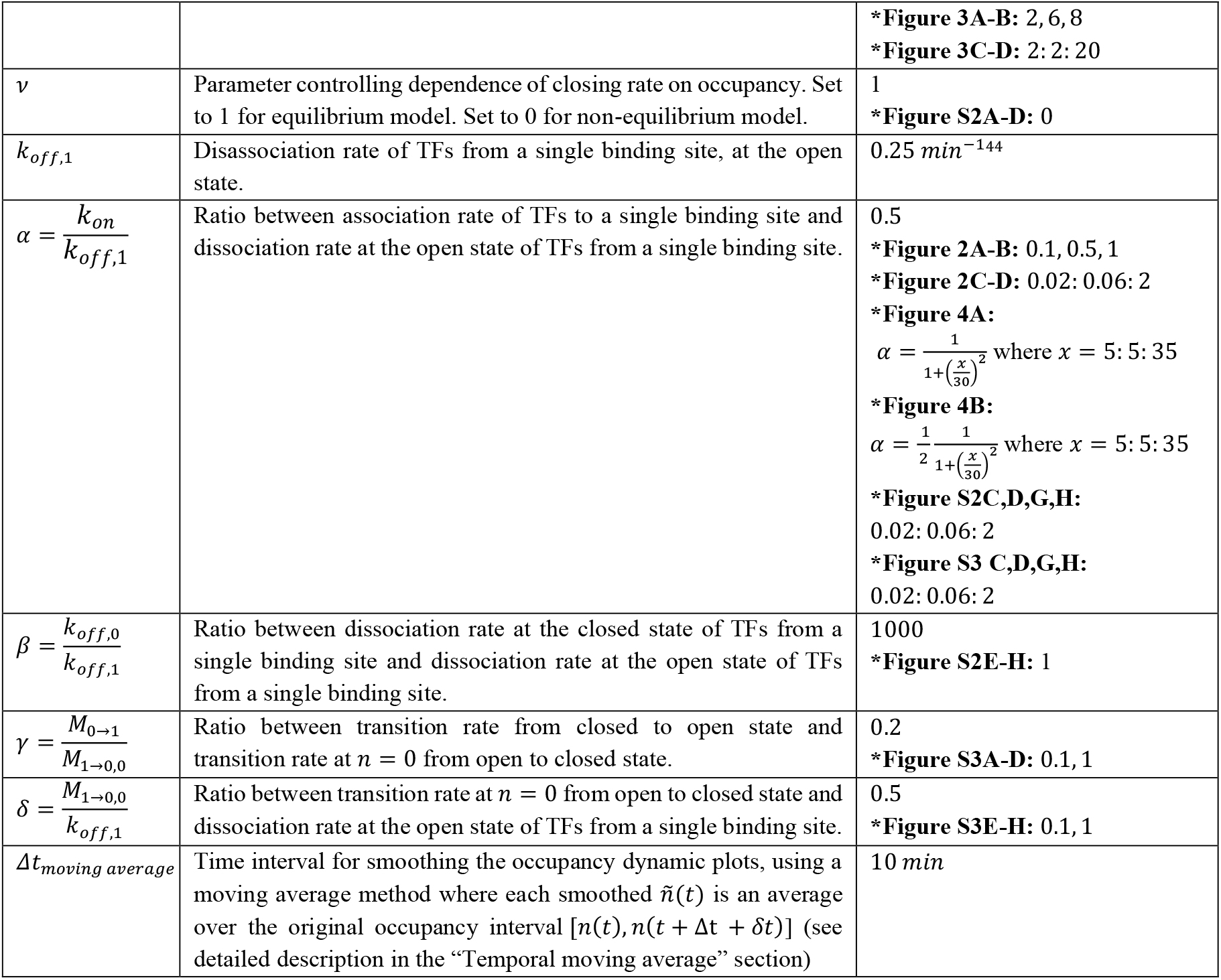
Parameters used to generate the results.

### Temporal moving average

The output of the simulation is an array of *n*(*t*). Since the temporal time intervals are non-uniform (each time step is different), we used a customized smoothing function which operates on a temporal window, rather than a finite amount of data points. This function calculates a smoothed value *ñ*(*t*) for each data point *n*(*t*), by averaging all data points in the interval [*n*(*t*), *n*(*t* + Δ*t* + *δt*)], where Δt is essentially the temporal window’s size, and *δt* accounts for small temporal gaps between *t* + Δt and its exact closest data point.

## Data and software Availability

Code generating the simulations data have been deposited in Github [Binshtok, 2023]. All other data are included in the manuscript and/or supporting information.

U. Binshtok, TSF *simulation* code, *GitHub*, https://github.com/Udi-Binshtok/TSF_model_2022, June 19, 2023

## Acknowledgments

We would like to thank ChangHwan Lee and Judith Kimble for providing the raw data of the experimental results in Fig. 4 (Lee et al.^12^.).

## Funding

DS and HGG were supported by Koret-UC Berkeley-Tel Aviv University Initiative in Computational Biology and Bioinformatics.

HGG was supported by the Winkler Scholar faculty Award in the Biological Sciences, by NIH Award R35GM158200, and by the Chan Zuckerberg Initiative Grant CZIF2024-010479. HGG is also a Chan Zuckerberg Biohub–San Francisco Investigator.

DS and BG were supported by an NSF/BSF award (2114950) and BG was supported by an NIH R01 award (GM079428).

## Author contributions

OA, UB, and DS performed calculations and simulations. OA, UB, DS, HGG, and BG wrote the manuscript.

## Competing interests

OA, UB, and DS declare no competing interests.

## SUPPORTING INFORMATION

### Supplementary figures

**Figure S1:**
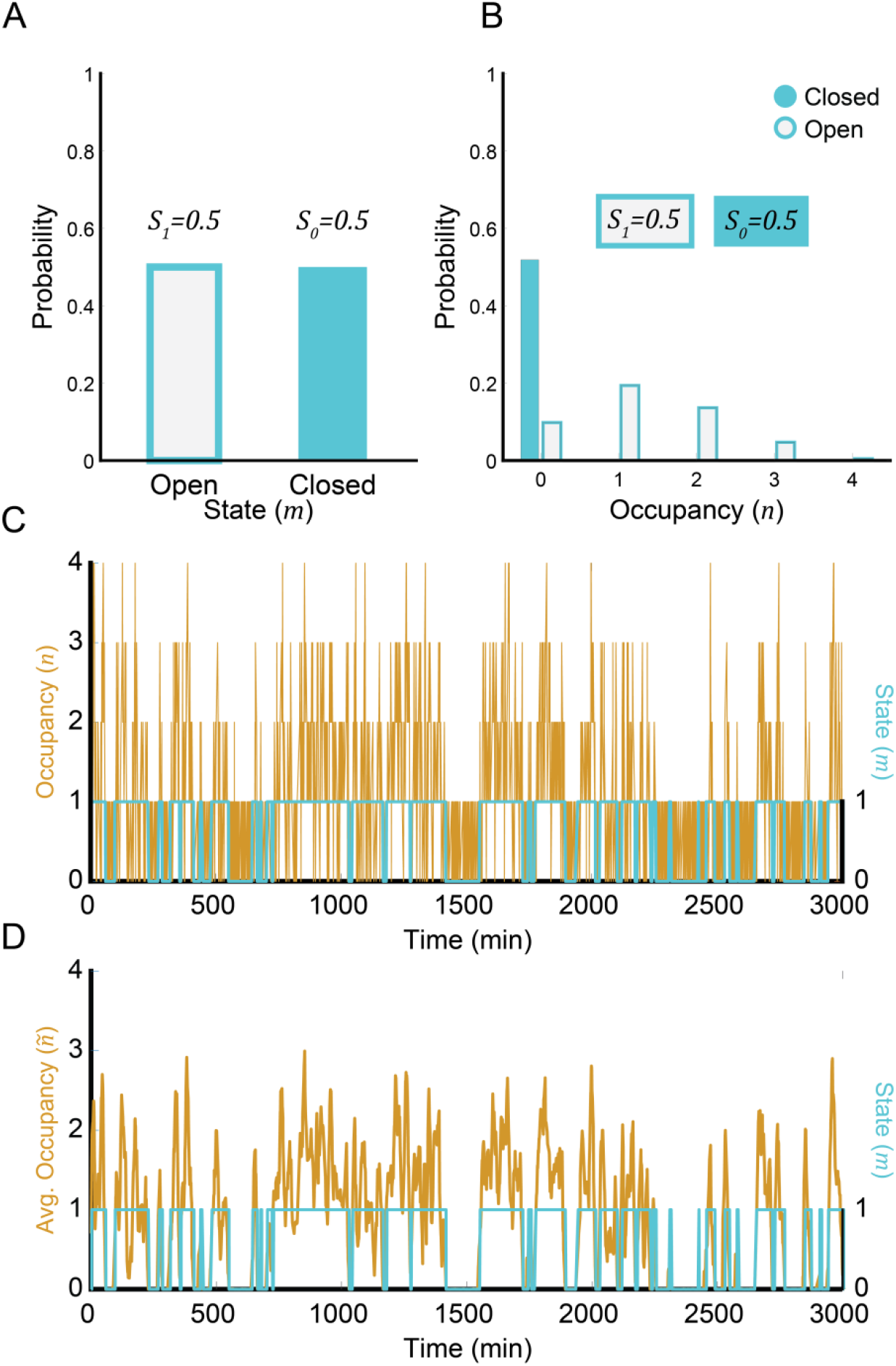
Comparison of raw occupancy, *n* (non-smoothened) and time averaged occupancy, *ñ* (smoothened) in simulations. **(A)** Probabilities of the system to be in the Open state (***S***_**1**_) or Closed state (***S***_**0**_), obtained from the analytical solution at steady state. **(B)** Probability distributions of binding site occupancy, calculated from the Gillespie simulation. The probability for each stat (Colored boxes above the bars) matches the distribution obtained from the analytic steady state solution. The probabilities are calculated from the full smoothed simulation (***t*** ∈ **[0, 30000**] ***min***; see also panel D). Default parameters were used (Supplementary Table 1). **(C)** Simulation of the raw (non-smoothed) occupancy, ***n***, using the default parameters (See Supplementary Table 1). **(D)** Time-averaged occupancy, ***ñ***. The time-averaged ***ñ*** is obtained by averaging the raw occupancy (shown in C) over a time interval of **Δt = 10 *min*** (See Methods and Supplementary Table 1).

**Figure S2:**
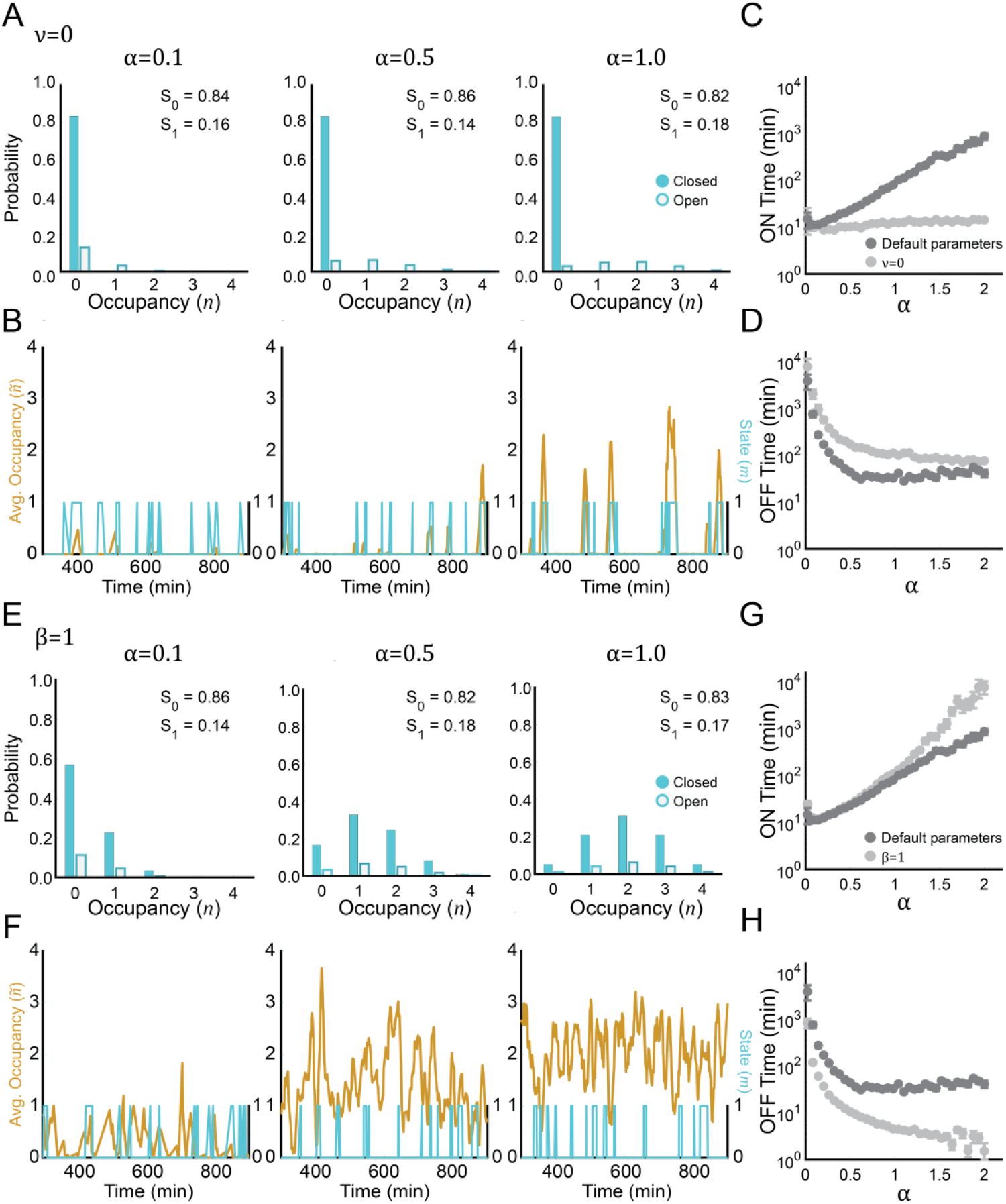
The TSF model in limits where no feedback is included in the model. **(A-B)** Simulations of a model with ***v* = 0**. In this model the transition rate from an open to closed state, ***M***_**1→0**_, is fixed and does not depend on the occupancy, ***n***. Hence, the probabilities of the system to be in the closed and open states, ***S***_**0**_ and ***S***_**1**_ respectively, do not change within numerical error when ***α*** changes (A). Accordingly, bursting dynamics do not depend strongly on ***α*** (B). **(C-D)** ON times and OFF times for different ***α*** values, calculated from simulations of the ***v* = 0** model (blue data points) and the TSF model with default parameters (Grey data points; same as in Fig. 2C-D). ON times remain relatively constant as ***α*** is varied, and OFF times saturate at higher values (compared to the TSF model). **(E-F)** Simulations of a model with ***β* = 1**. In this model the unbinding rate in the open state, ***k***_***off***,**1**_ is the same as the unbinding rate in the closed state, ***k***_***off***,**0**_. Hence, the occupancy distributions in the two states have a similar shape, and the probabilities of the system to be in the closed and open states, ***S***_**0**_ and ***S***_**1**_ (respectively), do not change when ***α*** changes (E). In this model the baseline level of occupancy increases with ***α***, but exhibits a noisy behavior rather than bursts (F). **(G-H)** ON times and OFF times for different ***α*** values, calculated from simulations of the ***β* = 1** model (blue data points) and the TSF model with default parameters (Grey data points; same as in Fig. 2C-D). ON times increase faster and OFF times decreasxe faster and to a lower value compared to the default model. In panels (C,D,G and H), each data point is an average ON time and average OFF time, calculated by analysis of the entire 30,000 minutes simulation and taking the average of continuous time periods when ***ñ* ≥ 1** (ON) and average of continuous time periods when ***ñ*** < **1** (OFF). Error bars are the standard error of the mean (SEM). In all panels, default parameters were used except those specified in the figure (See also Supplementary Table 1).

**Figure S3:**
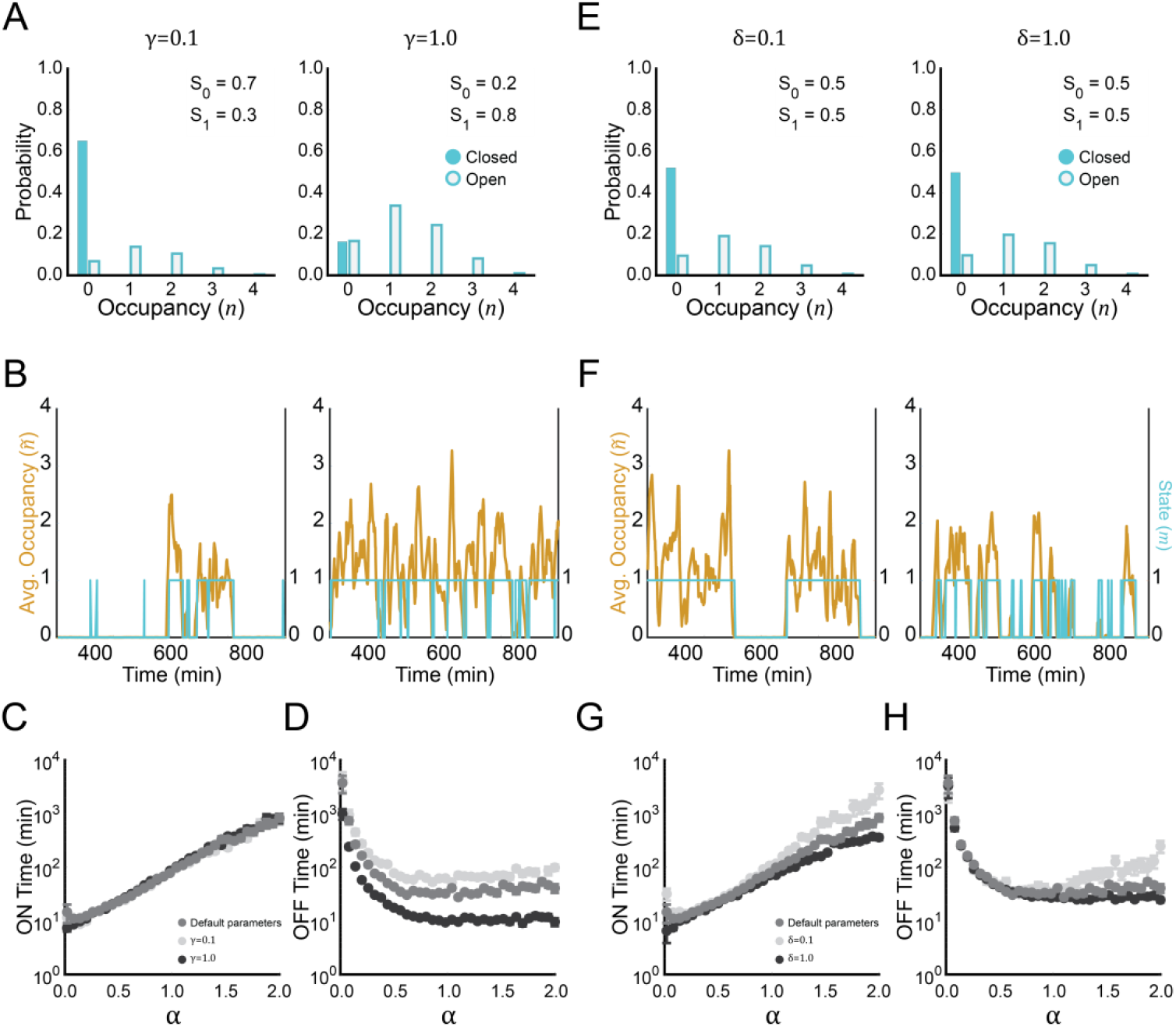
The TSF model is robust to changes in chromatin parameters *γ* or *δ*. **(A-D)** Dependence of occupancy distribution (A), dynamics (B), ON times (C) and OFF times (D) on the parameter 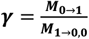. The effect of higher ***γ*** values is mainly to reduce the baseline value of the OFF times as seen in (D). **(E-H)** Dependence of occupancy distribution (E), dynamics (F), ON time (G) and OFF time (H) on the parameter 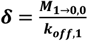. Minor quantitative differences are observed in bursting dynamics, that are more pronounced at high values of ***α***. In panels (C, D, G and H), each data point is an average ON time and average OFF time, calculated by analysis of the entire 30,000 minutes simulation and taking the average of continuous time periods when ***ñ* ≥ 1** (ON) and average of continuous time periods when ***ñ*** < **1** (OFF). Error bars are the standard error of the mean (SEM). In all panels, default parameters were used except those specified in the figure (Supplementary Table 1).

## Appendix

In this appendix we provide detailed derivation of Equation (9).

We start with the detailed balance equation

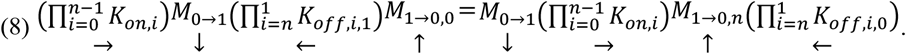

As 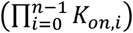 and *M*_0→1_ do not depend on the chromatin state or the occupancy, we can omit these expressions from both sides of the equation:

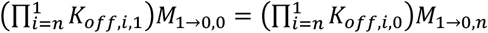

By dividing the equation by the TF disengagement term 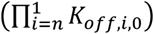, we can isolate the transition rate to the inactive chromatin state, *M*_1→0,*n*_:

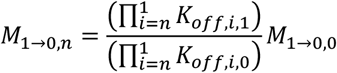

Plugging in the relation in equation (3), *K*_*off,n,m*_ = *nk*_*off,m*_, we get

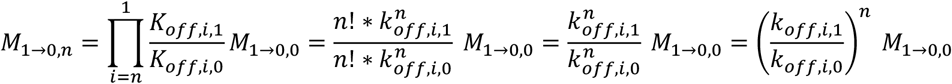

Lastly, since we defined

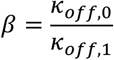

We get the following relationship:

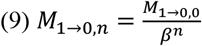

## Notes

### Competing Interest Statement

The authors have declared no competing interest.

